# ADHD symptoms map onto noise-driven structure-function decoupling between hub and peripheral brain regions

**DOI:** 10.1101/606228

**Authors:** Luke J. Hearne, Hsiang-Yuan Lin, Paula Sanz-Leon, Wen-Yih Isaac Tseng, Susan Shur-Fen Gau, James A. Roberts, Luca Cocchi

**Affiliations:** Clinical Brain Networks Group, QIMR Berghofer Medical Research Institute, Brisbane, Queensland, Australia; Department of Psychiatry, National Taiwan University Hospital, and College of Medicine, Taipei, Taiwan; Brain Modelling Group, QIMR Berghofer Medical Research Institute, Brisbane, Queensland, Australia; Institute of Medical Device and Imaging, National Taiwan University College of Medicine, Taipei, Taiwan; Graduate Institute of Brain and Mind Sciences, National Taiwan University College of Medicine, Taipei, Taiwan

## Abstract

Adults with childhood-onset attention-deficit hyperactivity disorder (ADHD) show altered whole-brain connectivity. However, the relationship between structural and functional brain abnormalities, the implications for the development of life-long debilitating symptoms, and the underlying mechanisms remain uncharted. We recruited a unique sample of 80 medication-naive adults with a clinical diagnosis of childhood-onset ADHD without psychiatric comorbidities, and 123 age-, sex-, and intelligence-matched healthy controls. Structural and functional connectivity matrices were derived from diffusion spectrum imaging and multi-echo resting-state functional MRI data. Hub, feeder, and local connections were defined using diffusion data. Individual-level measures of structural connectivity and structure-function coupling were used to contrast groups and link behavior to brain abnormalities. Computational modeling was used to test possible neural mechanisms underpinning observed group differences in the structure-function coupling. Structural connectivity did not significantly differ between groups but, relative to controls, ADHD showed a reduction in structure-function coupling in feeder connections linking hubs with peripheral regions. This abnormality involved connections linking fronto-parietal control systems with sensory networks. Crucially, lower structure-function coupling was associated with higher ADHD symptoms. Results from our computational model further suggest that the observed structure-function decoupling in ADHD is driven by heterogeneity in neural noise variability across brain regions. By highlighting a neural cause of a clinically meaningful breakdown in the structure-function relationship, our work provides novel information on the nature of chronic ADHD. The current results encourage future work assessing the genetic and neurobiological underpinnings of neural noise in ADHD, particularly in brain regions encompassed by fronto-parietal systems.

## Introduction

Adult attention-deficit hyperactivity disorder (ADHD) is a common neurodevelopmental disorder characterized by inattentive and hyperactive-impulsive symptoms beginning in early childhood [1]. Identifying the neural underpinnings of adult ADHD is an ongoing research endeavor, critical to the definition of neural mechanisms supporting clinical outcomes of childhood-onset ADHD and the development of novel targeted interventions [2].

Neuroimaging work has provided important insights into altered structural [3–5] and functional [6,7] brain connectivity underpinning ADHD pathophysiology, and suggest that network interactions, rather than regional abnormalities, contribute to phenotypic expression of the disorder [8]. Anatomically, results have been mixed. Recent studies have shown no changes in the ADHD connectome [9], whereas others have pointed to various abnormalities in white matter tracts including the corpus callosum and posterior circuits related to the limbic and occipital systems, the fronto-striato-cerebellar connections, and pathways linking default-mode and fronto-parietal hub regions [4,5,10].

Complementing findings from diffusion MRI, resting-state functional magnetic resonance image (rs-fMRI) studies have highlighted that both diagnosis and symptoms of ADHD are linked to reduced segregation between the activity of control networks supporting external task engagement and the default-mode network [6,7,11]. Reduced functional connectivity within, and between, the default-mode, sensory, and control networks has also been reported both in children and adults with ADHD [6,7,10,11].

Emerging evidence suggests that patterns of functional connectivity are constrained by their anatomical underpinning: The connectome [12,13]. Structural and functional brain network alterations in adult ADHD partially overlap [10], but the direct link between these structure-function aberrations has not been formally explored. A candidate mechanism for altered structure-function associations is excessive neural noise: The increased random variability in neural activity [14–16]. Evidence for this idea comes from several related lines of research at different levels of description. Variable and inconsistent behavior, like those observed in ADHD [17,18] and other contexts [e.g., learning, aging, developmental dyslexia 19–21], has been suggested to correlate with increased neural noise. A number of neuroimaging studies have also highlighted increased brain signal variability in ADHD [22–25]. At the neuronal and molecular levels, drug treatments that are effective in ADHD by targeting catecholaminergic pathways are thought to modulate neural signal-to-noise ratios [26–30].

Here, we used multi-echo rs-fMRI and diffusion spectrum imaging (DSI) to investigate possible changes in whole-brain structure-function coupling in a large sample of well-characterized, medication-naïve adults with childhood-onset ADHD and matched healthy controls [11]. Based on previous findings [11] and the hypothesis that psychiatric conditions are primarily pathologies of brain hubs [31], we expected significant departures from the typical structure-function coupling in ADHD. Specifically, a breakdown in the structure-function association is likely to occur in connections involving brain hubs that belong to the control and default-mode brain networks [31,32]. To investigate neural noise as a possible mechanism of this structure-function decoupling [33], we adopted whole-brain computational modeling. Our model explicitly tested the hypothesis that increased heteroscedasticity in the levels of intrinsic neural noise drives the expected breakdown in the structure-function coupling. Heteroscedasticity occurs when the variance of explanatory variables – neural noise level – is not identical across brain regions.

## Methods

### Sample

We recruited 80 psychotropic-nai□ve adults with childhood-onset ADHD aged 18–39 years (mean 26.7 years), who fulfilled DSM-IV-TR criteria for the current diagnosis of ADHD. While this cohort may be narrow in terms of typical clinical ADHD phenotypes, our carefully selected sample allowed the unequivocal assessment of ADHD-specific structural and functional brain networks in the absence of common confounds including other neurodevelopmental disabilities, psychotropic exposure and major psychiatric comorbidities [10]. Results from the clinical sample were benchmarked against the findings of 123 age-(mean 25.7 years), sex-, and IQ-matched healthy controls. Participants were assessed at the Department of Psychiatry, National Taiwan University Hospital (NTUH), Taipei, Taiwan. Details regarding the recruitment procedure are described elsewhere [11] (**Supplementary Methods**).

### MRI acquisition and preprocessing

Brain imaging data were acquired with a Siemens 3T Tim Trio scanner equipped with a 32-channel head coil. Details regarding the preprocessing and quality control of the multi-echo resting-state and diffusion data are described in the **Supplementary Methods (Supplementary Figure 1)**. The final sample included 78 ADHD adults and 118 healthy controls (**Table 1**).

**Table 1.** Demographic and clinical features of the participants.

| Mean (SD) | Control (N=118) | ADHD (N=78) | Statistics |
| --- | --- | --- | --- |
| **Age (18-39 years)** | 25.8 (5.0) | 26.6 (5.5) | $p = 0.287$ |
| **Sex (M/F)** | 76/42 | 54/24 | $p = 0.484$ |
| **FIQ** | 109.8 (9.3)<br>(range: 89-138) | 107.5 (10.4)<br>(range: 80-137) | $p = 0.101$ |
| **VIQ** | 108.2 (9.0) | 105.7 (11.2) | $p = 0.088$ |
| **PIQ** | 110.4 (11.4) | 108.3 (16.3) | $p = 0.289$ |
| **ADHD symptoms** |  |  |  |
| **SNAP-IV (Parent-report)<sup>a</sup>** |  |  |  |
| **Inattention (0-27)** | 6.6 (4.9) | 19.6 (5.0) | $p < 0.001$ |
| **Hyperactivity/Impulsivity (0-27)** | 3.2 (4.4) | 13.4 (6.4) | $p < 0.001$ |
| **ASRS (Self-report)** |  |  |  |
| **Inattention (0-36)** | 13.3 (5.2) | 27.0 (4.8) | $p < 0.001$ |
| **Hyperactivity/Impulsivity (0-36)** | 9.1 (5.2) | 19.9 (6.3) | $p < 0.001$ |
| **Mean frame-wise displacement<sup>b</sup> (mm)** | 0.045 (0.021)<br>(range: 0.014-0.123) | 0.048 (0.024)<br>(range: 0.017-0.108) | $p = 0.354$ |
| **Signal dropout counts<sup>c</sup>** | 30.8 (22.4) | 28.8 (21.4) | $p = 0.536$ |

<sup>a</sup> Measured by the parent-rated Swanson, Nolan, and Pelham, version IV (SNAP-IV) scale.
<sup>b</sup> A summary estimate of in-scanner motion levels of resting-state fMRI, as estimated by the Euclidian norm (enorm: square root of the sum of squares of the differences in motion derivatives), computed with AFNI's 1d\_tool.py.
<sup>c</sup> A summary estimate of in-scanner motion levels of diffusion spectrum imaging (see the Methods).
Abbreviation: ADHD=attention-deficit hyperactivity disorder; FIQ=full intelligence quotient; PIQ=performance intelligence quotient; VIQ=verbal intelligence quotient; ASRS=Adult ADHD Self-Report Scale; M=male; F=female; R=right; L=left; SD=standard deviation.

### Structural and functional brain network construction

We generated whole-brain structural (SC) and functional (FC) connectivity matrices for each individual, based on a common and recently validated cortical parcellation [34] (**Fig. 1A**, see **Supplementary Information** for control analyses). Fourteen additional subcortical structures from the Harvard-Oxford atlas were added to the parcellation, resulting in 214 total regions (*Schaefer-214* henceforth; **Supplementary Table 1**). Individual whole-brain tractography maps were combined with the pre-defined anatomical boundaries defined by this *Schaefer-214* parcellation to generate a weighted SC matrix (**Fig. 1B**). Each edge of the network corresponds to the total number of normalized streamlines that interconnect any two brain regions, adjusted for the interregional fiber length [35]. For resting-state data, regional time-series were calculated as the mean across voxels within each region included in the brain parcellation. For each individual, Pearson’s correlations were calculated between the time-series of all regions to calculate FC. Finally, a Fisher z-transformation was applied to the FC matrices.

**Fig. 1.**
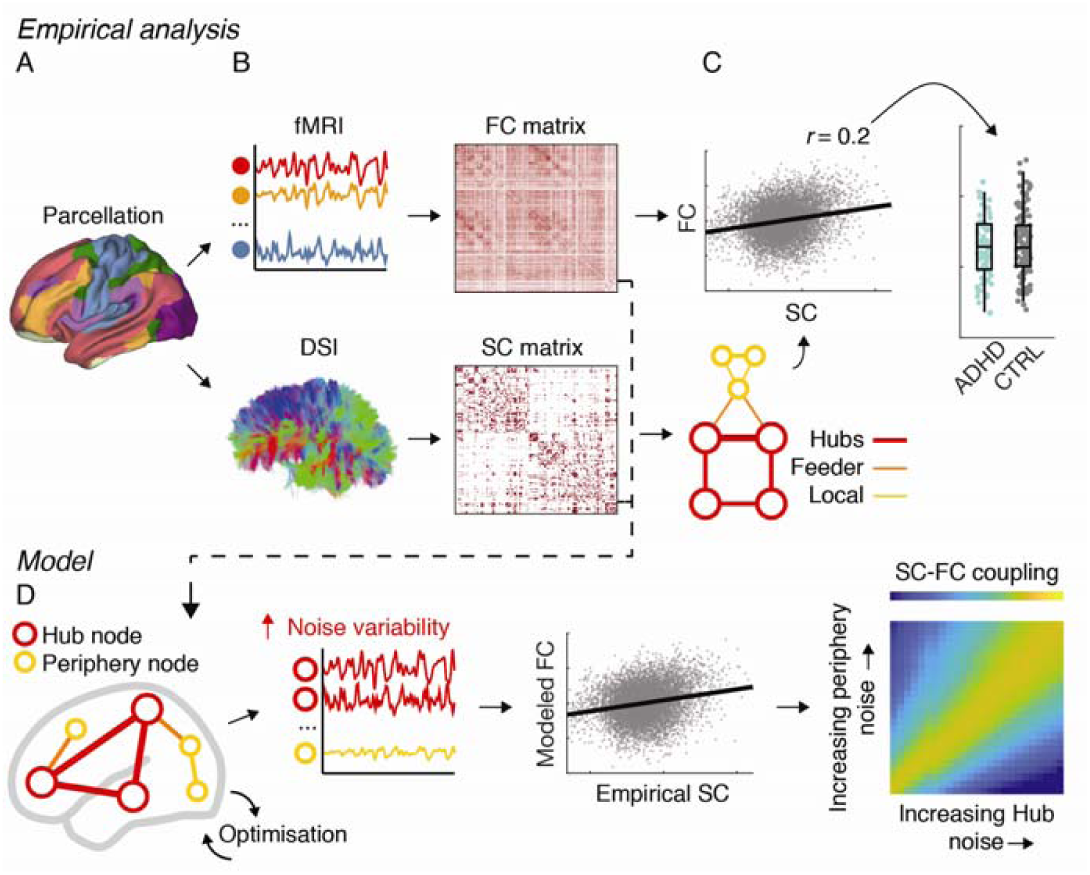
Conceptual overview of the analysis pipeline. **A.** Analyses were conducted using a whole-brain parcellation including 214 cortical and subcortical regions. Replication analyses were performed using two alternative brain parcellations (see text). **B.** Structural (SC) and functional connectivity (FC) matrices were derived from diffusion spectrum imaging (DSI) and multi-echo resting-state fMRI data, respectively. Darker colors indicate higher normalized streamline counts (SC) and higher Fisher-z normalized Pearson’s correlation values between every possible pair of brain regions (FC). **C.** The topological organization of the SC matrices was examined to derive measures of different connection types: hub connections, feeder connections, and local connections. Individual-level correlations between SC and FC were used to estimate structure-function coupling, which was then analyzed with between-group statistics. **D.** A computational model was used to assess the potential neural mechanisms that lead to decreased structure-function coupling. Empirical SC was used as input in the model and model parameters were estimated by fitting to empirical FC. We systematically assessed if an increase in the noise heterogeneity in hub or peripheral nodes could result in a marked dissociation between functional and structural connectivity.

### Connection classes

We identified hub regions according to an aggregate ranking across multiple metrics including degree, strength, subgraph centrality, and betweenness [36,37]. The top 15% composite scores (N = 32, **Supplementary Table 1&2, Supplementary Figure 2**) were used to identify hub regions within each individual; all other nodes were assigned as *periphery* nodes. Hub *connections* were defined as edges that connected any two hub nodes. Feeder connections linked hub nodes to periphery nodes, and local connections linked periphery nodes (**Fig. 1C**) [32,38].

### Structure-function relationships

Brain network structure-function relationships were conducted in line with previous research [32]. First, non-zero SC values within each individual connectome were isolated and normalized using a rank-based inverse Gaussian transformation [39]. The resulting SC values were correlated with corresponding FC values (i.e., the same edges). This analysis produced a single Pearson’s *r* value that summarized the global structure-function association for each individual [40]. These values were used to populate group distributions and were subsequently contrasted using between-group statistics. This entire procedure was completed at the level of the whole network and within each respective connection class: hubs, feeders, and local edges.

Previous work investigating resting-state networks, including data from the current cohort [11], has highlighted the key role of control, default-mode, and sensory networks in adult ADHD [6,7]. Based on these results, we also tested for specific changes in SC-FC coupling within these networks. A minimum of 50 of edges was used to infer structure-function relationship, thus control networks were defined as the combination of fronto-parietal, alongside dorsal and ventral attention affiliations from the adopted parcellation, while sensory connections included both visual and somatomotor affiliations. Default-mode connections were as in the original parcellation. Once SC-FC coupling was estimated within each network, the mean *r* values (Control-ADHD) were presented within and between each network.

### Relationship between structure-function coupling and behavioral symptoms of ADHD

Given the notion that measures of ADHD symptoms are continuously distributed in the general population [41,42], we investigated brain-behavior relationships across both ADHD and control groups (**Fig. 1C**). Inattention and hyperactivity-impulsivity symptoms based on the parent-rated Swanson, Nolan, and Pelham, IV (SNAP-IV) [43] and self-rated Adult ADHD Self-Report Scale (ASRS) [44] (**Table 1**) were used in the analysis. These four symptom items (two from each measure) were transformed using a rank-based inverse Gaussian, then entered into a principal component analysis to reduce the dimensionality of the data. The first component, accounting for 81% of the variance, was then correlated with the structure-function coupling of the whole sample (**Supplementary Table 3**).

### Statistical comparisons between groups

To ensure that the SC density did not explain between-group differences, summed binary and weighted degrees were compared between groups. Average connection weights within each *connection class* were compared between each group. In addition, the network based statistic (NBS) [45] was used to explore any possible differences in SC between controls and ADHD (5000 permutations, threshold *t* = 3). ADHD-associated alterations of FC using NBS have been reported in our initial study on this sample [11].

Mann–Whitney U tests were used to identify possible differences in the structure-function association between control and ADHD groups. Bonferroni correction (family-wise error rate, FWE) for multiple comparisons was applied to follow-up statistics, with α_FWE_ < 0.05 indicating statistical significance. Effect sizes (*r*_*eff*_ were reported for all tests using the formula 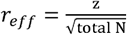 (Rosenthal, 1994). For this metric, *r*_*eff*_ = 0.1 is considered a small effect, *r*_*eff*_ = 0.3 is considered medium, and *r*_*eff*_ = 0.5 is considered a large effect [46]. Statistical analyses were performed in MATLAB (Mathworks) with code available online (https://github.com/ljhearne/ADHDSCFC).

### Computational modeling: Assessing the neural factors driving structure-function breakdown

We adopted whole brain computational modeling to simulate SC-FC coupling. We tested the hypothesis that increased heteroscedasticity in the levels of intrinsic neural noise was associated with differences in SC-FC coupling between groups. Specifically, we manipulated the levels of heteroscedasticity, which occurs when the variance of explanatory variables (i.e., neural noise level) is not identical across brain regions. The model incorporates SC to represent the strength of connections between brain regions. In addition to the weights specified in the empirical SC matrix, structural connections are scaled by a global coupling parameter. This parameter can then be varied systematically to simulate and compare the global dynamics emerging from the model with the empirical FC derived from the rs-fMRI data.

We chose a simple stochastic linear model of the Ornstein-Uhlenbeck type [47–49]. The main motivations behind this choice were that the model: (i) allows us to simulate whole-brain patterns of FC *from* SC matrices; (ii) enables tests of the hypothesis that increased heteroscedasticity of neural noise levels results in a breakdown in structure-function coupling; (iii) can be considered a generic linearization of more complex models with a stable fixed point (a mathematical approach at the core of e.g. dynamic causal modeling for fMRI [50]); and (iv) permits a direct analytical derivation of FC from empirical SC without the need of computationally demanding numerical simulations **(Supplementary Figure 3&4)**. The model equation is:

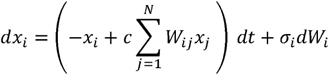

where *x*_*i*_ is the activity of the *i*-th region; *c* is the global coupling strength which rescales the strength of structural connections of the system; *W*_*ij*_ is the connectivity weight to region *I* from region *j* (as specified by the empirical SC matrix); *σ*_*i*_ is the intrinsic noise amplitude/level of the *i*-th region, and defines the size of random increments *σ*_*i*_*dW*_*i*_ in the dynamics of the region, and *N* is the total number of regions in the connectome. Previous modeling studies [48,49] have considered the noise levels to be constant across the whole network (i.e., all *σ*_*i*_ are identical). In light of previous suggestions [14,51–53], we hypothesized that heteroscedasticity across a specific subset of brain regions (hubs or periphery) would have a detrimental impact on SC-FC decoupling. To test our hypothesis, we systematically analyzed varying degrees of heteroscedasticity in the noise levels in distinct subsets of regions independently (hub and periphery regions). A comprehensive description of the modeling can be found in the **Supplementary Methods (Supplementary Figure 5)**.

## Results

### Similar structural connectivity between groups

Results showed no difference in weighted (*p* = 0.89, *z* = 0.13, *r*_*eff*_ = 0.01), or unweighted (*p* = 0.24, *z* = −1.19, *r*_*eff*_ = −0.08) summed degree across groups. Likewise, the whole-brain network-based statistics comparing ADHD and healthy control groups revealed no significant differences in structural connectivity between the groups (ADHD > controls, *p* = 0.63; controls > ADHD, *p* = 0.78). Next, we sought to investigate potential differences in *classes* of structural connections, namely hubs, feeders, and local connections. No significant group differences were observed when comparing mean connection strength within hub (*p* = 0.86, *z* = −0.17, *r*_*eff*_ = −0.01), feeder (*p* = 0.77, *z* = −0.29, *r*_*eff*_ = 0.04), or local connections (*p* = 0.23, *z* = 1.21, *r*_*eff*_ = 0.09).

### Structure and function coupling in ADHD is reduced in feeder connections

When considering all edges within the network, results indicated a significant difference in SC-FC coupling (p = 0.01, z = 2.51, r_eff_ = 0.18, **Fig. 2A**). We then assessed the contribution to this effect of each connection class (hub, feeder or local). Results showed that compared to controls, ADHD had a significantly lower SC-FC association in feeder connections of a non-trivial effect size (p_FWE =_ 0.005, z = 3.10, r_eff_ = 0.22). No between group differences were found in hub (p_FWE_ = 1, z = 0.55, r_eff_ = 0.04) or local (p_FWE_ = 0.33, z = 1.60, r_eff_ = 0.11) connections (**Fig. 2A**).

**Fig. 2.**
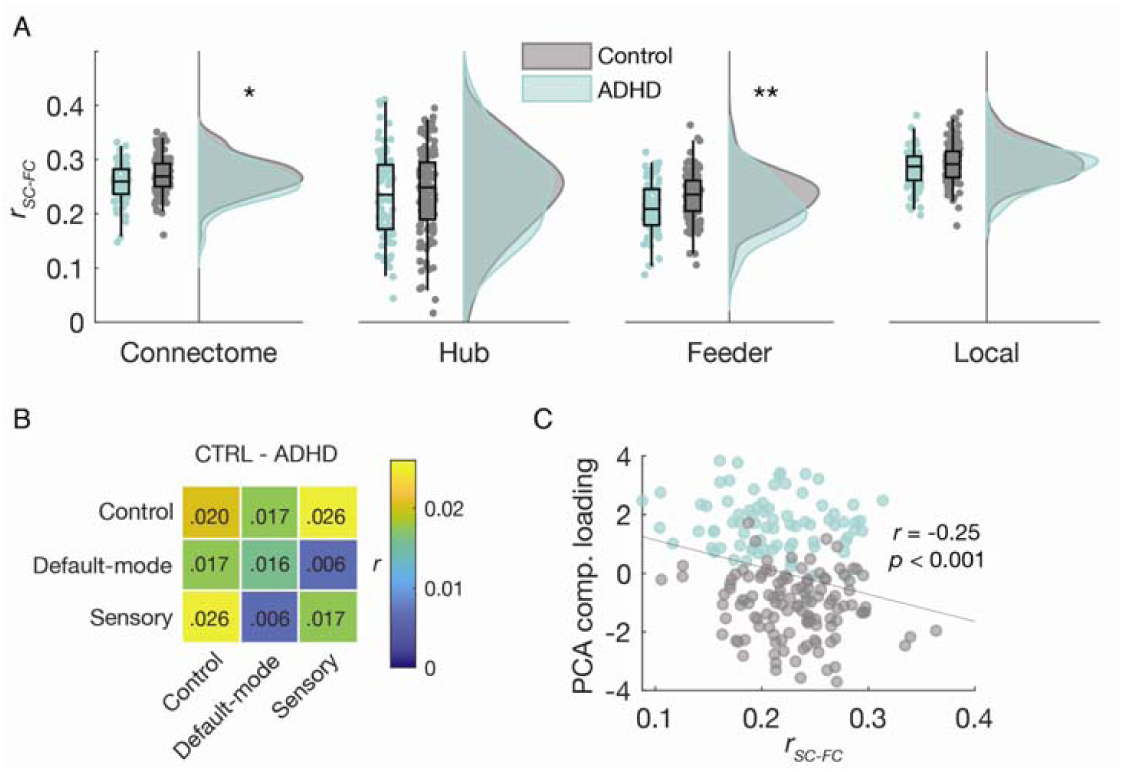
Structure-function relationships in drug-naïve adults with ADHD and healthy matched controls. **A.** Distributions of *r* values across the whole connectome and the three connection classes[73]. Significant differences between ADHD and Control groups were observed in the whole connectome but were driven by a large group difference in feeder connections. **B.** Mean differences in SC-FC coupling (Controls *minus* ADHD) when constrained to feeder connections within and between control, default-mode, and sensory functional networks. The largest deficit in SC-FC coupling in ADHD compared to controls were found between control and sensory network connections (*r* = 0.026). **C.** Correlation between symptoms and SC-FC coupling in feeder connections. SC-FC coupling strength was negatively correlated with the ADHD symptom factor scores derived from principal components analysis. * < 0.05, ** < 0.01 corrected for multiple comparisons.

### Feeder structure-function decoupling in control, default-mode, and sensory brain networks

To further explore the anatomical specificity of the observed deficits in structure-function coupling, we isolated feeder connections that belonged to control, default-mode, or sensory (merging somatomotor and visual) networks. As per the previous analysis, we correlated SC and FC values for connections within and between the selected brain networks. This resulted in a three-by-three matrix for both ADHD and healthy control groups that represented the degree of SC-FC coupling within and between control, default mode, and sensory networks. The largest reduction in SC-FC associations in ADHD compared to healthy controls were located in connections between control and sensory networks (**Fig. 2B**).

### The magnitude of structure-function decoupling correlates with the severity of ADHD symptoms

Individual symptom scores captured by PCA linearly correlated with indices of structure-function coupling in feeder connections, such that lower structure-function coupling was associated with more severe ADHD symptoms (*p* = 0.0004, *r* = −0.25, **Fig. 2C**).

### Noise in hubs and periphery as a neural mechanism for structure-function breakdown

Finally, we sought a neural mechanism for how altered structure-function relationships could emerge in the absence of significant differences in the connectome. In particular, we aimed to use computational modeling to explain our finding of selective deficits in feeder connection SC-FC coupling. We systematically explored two scenarios with noise heteroscedasticity – i.e., increased heterogeneity in the intrinsic neural noise levels *σ*_*i*_ across brain regions.

In the first scenario, we analyzed the case of heterogeneity between hubs and periphery (*σ*_*H*_ ≠ *σ*_*P*_) for hub nodes (*H*) and peripheral regions (*P*), maintaining *σ*_*H*_ and *σ*_*P*_ constant *within* each class of regions. Exploring ranges of *σ*_*H*_ and *σ*_*P*_ (**Fig. 3A-C**) we analyzed the changes in SC-FC coupling for the three classes of connections (hub, feeder, and local). We found that feeder connections were the most susceptible to subtle imbalances between intrinsic noise levels in hub and periphery regions, reflected in the quick decrease in SC-FC coupling (**Fig. 3B**). On the contrary, hub and local connections exhibited only small changes (**Fig. 3A&C**). Specifically, a small imbalance such that *σ*_*H*_ < *σ*_*P*_, with *σ*_*P*_ 10% larger than *σ*_*H*_, produced a slight (< 2%) reduction in SC-FC coupling in hubs compared to the homogenous *σ*_*H*_ = *σ*_*P*_ case, similar to the empirically observed slight decrease for hub connections in **Fig. 2A** (< 2%). Conversely, a 10% imbalance in the opposite direction (*σ*_*H*_ > *σ*_*P*_) yielded a negligible (∼0.3%) increase in hub SC-FC coupling. The increased sensitivity of feeder connections was demonstrated by the same 10% imbalance (*σ*_*H*_ < *σ*_*P*_) resulting in a 4% decrease in SC-FC coupling for feeder connections compared to the homogenous case. Importantly, an imbalance of approximately 50% (*σ*_*H*_ < *σ*_*P*_) was required to obtain the 10% decrease in SC-FC coupling empirically observed in ADHD feeder connections (**Fig. 2A**). This larger imbalance also resulted in a < 2% reduced SC-FC coupling in hub connections, again in accordance with empirical results. Thus, larger differences between mean noise amplitude levels in hubs and periphery led to greater SC-FC decoupling specific to feeder connections, mirroring the selective deficits observed in ADHD.

**Fig. 3.**
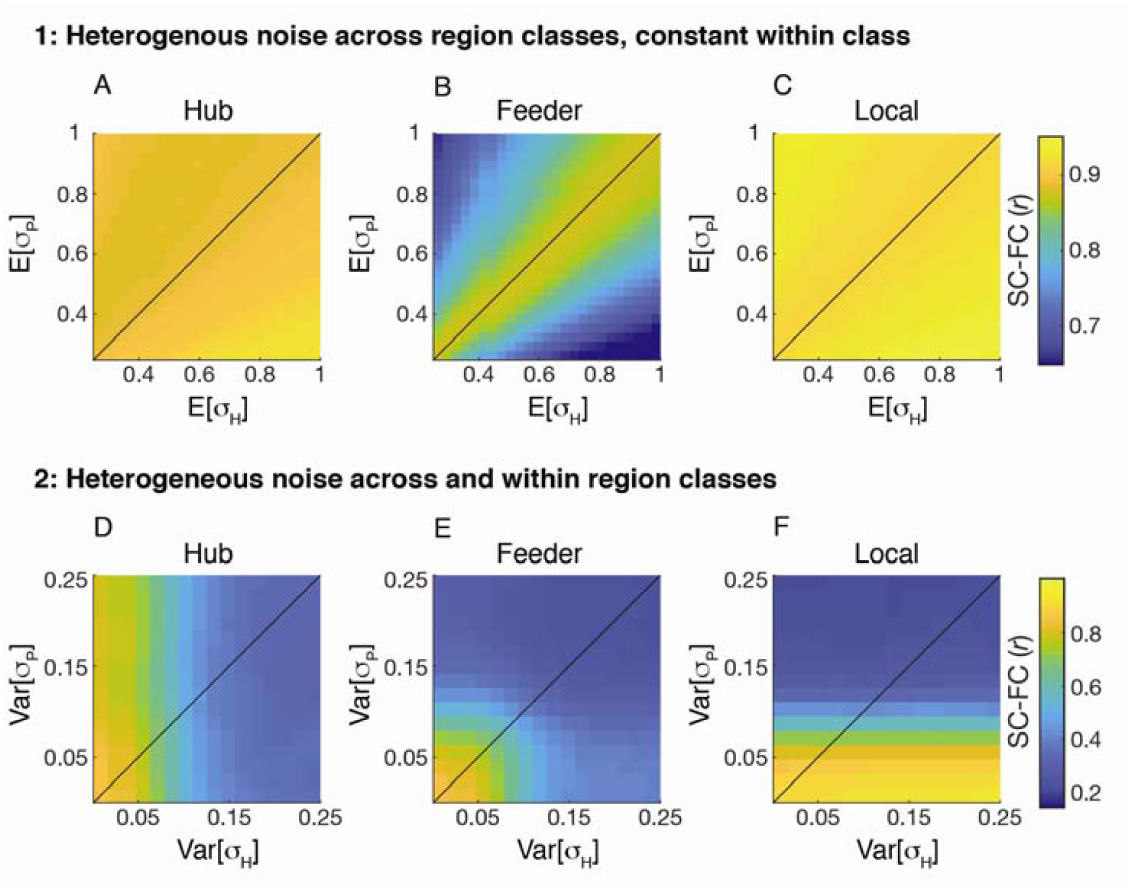
Modeling the effect of noise heteroscedasticity on structure-function coupling. Effects of noise heteroscedasticity on SC-FC coupling. *Top row*: Scenario 1 - Noise heterogeneity between hubs and periphery (*σ*_*H*_ ≠ *σ*_*P*_) for hubs (H) and peripheral brain regions (P), *σ*_*H*_ and *σ*_*P*_ constant within each class of regions (hubs and periphery). *Bottom row*: Scenario 2 - noise levels (*σ*_*i*_) within hubs and periphery varied from region to region. The colormaps quantify the SC-FC coupling (Pearson correlation between SC and FC matrix entries). **A/D**. Hub connections. **B/E**. Feeder connections. **C/F**.Local connections. E[·] = expected mean value; Var[·] = variance. The line in each panel corresponds to the case E[*σ*_*H*_] = E[*σ*_*P*_] (top row) or Var[*σ*_*H*_] = Var[*σ*_*P*_] (bottom row).

In the second scenario, we modeled the case where the noise levels (*σ*_*i*_) within hubs and periphery also varied from region to region. This allowed us to examine whether heteroscedasticity within hubs and/or periphery regions could contribute to the observed disruption of SC-FC coupling in ADHD. We systematically explored ranges of variance (Var[*σ*_*H*_] and Var[*σ*_*P*_]) fornoise levels normally distributed around means (E[*σ*_*H*_] and E[*σ*_*P*_]), set here such that E[*σ*_*P*_] is 10% larger than E[*σ*_*H*_] in line with the above results for hub connections (comparing **Fig. 3A** to **Fig. 2A**). We found that connections within a region class (i.e., hub-hub or periphery-periphery) are resilient to increased variability of intrinsic noise levels in the opposite type. Indeed, the SC-FC coupling in hub connections (**Fig. 3D**) and local connections (**Fig. 3F**) remained almost constant for increased noise variability in peripheral and hub regions, respectively. However, feeder connections (**Fig. 3E**) are clearly susceptible to changes in noise level heterogeneity within either hub or periphery regions, which implies an increased sensitivity to heteroscedasticity could also contribute to the disruption of SC-FC coupling in ADHD.

## Discussion

The present study provides evidence of a clinically significant breakdown in brain structure-function (SC-FC) coupling in medication-naive adults with childhood-onset ADHD. In line with the hypothesis that hub regions are critically vulnerable to brain pathology [31,32,54], ADHD was associated with a marked SC-FC decoupling in connections linking brain hubs to peripheral regions (feeders) within and between control and sensory networks. Modeling results further suggest that such decoupling is potentially linked to: (i) an imbalance in noise amplitudes in hubs and the periphery (e.g., increased ‘unreliability’ in signals originating from the periphery) and, (ii) higher peripheral heteroscedasticity (i.e., the peripheral noise is more diverse and more difficult for the hubs to filter out). Altogether, results from this work propose a novel neural mechanism explaining structure-function decoupling in brain connectivity underpinning the chronic manifestation of ADHD symptoms.

Structural networks are thought to place significant constraints on FC and local brain activity [12,33,40]. The decoupling between FC and its structural basis is therefore thought to represent a key index of brain network pathology in psychiatric illnesses including schizophrenia [32,55,56]. Our results are in line with the general notion that a structure-function breakdown in psychiatric illnesses involves anatomically defined hub brain regions [31]. The observed association with behavior, indicating that reduced structure-function coupling in feeder connections is related to higher severity of ADHD symptomology, provides support for the clinical relevance of this deficit in ADHD. By using a parsimonious model explaining the emergence of functional connectivity from underlying anatomical connectivity, we found that increased heteroscedasticity in intrinsic noise levels, either in hubs or periphery, has a strong detrimental effect in feeder connections, and to a lesser extent in hub-hub connections.

Physiologically, reduced SC-FC coupling due to increased noise heteroscedasticity in peripheral regions can be understood as brain hubs being unable to average out incoming peripheral functional disruptions. This adds weight to the notion that ADHD symptoms may arise from increased neural noise in the activity of frontal hub regions composing fronto-parietal and default-mode networks [23]. These brain networks have been tied to psychological functions critically impacted by ADHD, including cognitive control, sustained attention, and behavioral variability [57–60]; with activity being shown to be more variable in ADHD compared to controls [22–24]. This abnormal brain network activity can be, at least in part, restored by methylphenidate treatment [61–63]. In line with the above, a proposed mechanism for this therapeutic effect is the modulation of neural noise’s characteristic 1/f^α^ spectrum [15]. Because our model dynamics have a 1/f^α^ noise spectrum (**Supplementary Figure 4**), current results provide support for this hypothesis.

Our empirical findings showed that feeder connections are the most affected by the decoupling between function and anatomy. Feeder connections comprise long-range anatomical routes allowing efficient communication between remote brain regions belonging to different brain networks [38]. We here found that connections within control networks, as well as between regions comprising control and sensory networks, contributed to the overall reduction in structure-function association in ADHD. These findings are in agreement with previous neuroimaging studies in ADHD [6,7,64,65] and healthy controls [58,66], highlighting the key role of these connectivity patterns to support normal and pathological attention and inhibitory processes. We also note that altered patterns of FC, and SC-FC decoupling, can occur in the absence of deficits in SC [55]. In fact, whereas white matter connections are predictors of FC [40], the opposite is not always true [67].

The absence of significant group differences in the structural connectome is at odds with some previous reports [3,4]. Due to the sample size and the quality of the data, it is unlikely that the negative finding reported here is due to a lack of statistical power in detecting meaningful differences in the ADHD connectome. Moreover, our result is consistent with recent work showing the existence of FC abnormalities with preserved white matter properties in ADHD [68]. The discrepancy between our findings and earlier literature [3] may be explained by non-neural factors. For example, the absence of significant differences between the ADHD and control connectomes reported here may reflect our emphasis on comparable levels of head motion between the two groups; a critical factor that produce spurious group differences in ADHD [3,69]. Our cohort of psychotropic-naive adults with established childhood-onset ADHD in the absence of co-occurring psychiatric conditions may also contribute to this negative finding, as psychostimulant exposure [70] and comorbidity [71] have been reported to affect SC in ADHD. Whereas our results cannot completely exclude the presence of altered white matter integrity in ADHD, they suggest that any such differences are small overall, and the manifestation of core ADHD symptoms is underpinned by functional deregulations and related decoupling in SC-FC. Further work in broader clinically-representative samples will be necessary to parse the contributions of factors including comorbidities and medication to the integrity of the connectome [10,72].

By combining functional and diffusion-weighted imaging with computational modeling, our study has advanced the understanding of neural mechanisms that underpin chronic ADHD symptoms. More specifically, our work showed that a clinically meaningful function-structure decoupling in ADHD is likely related to increased neural noise heterogeneity between hubs and periphery regions. This knowledge is consistent with the positive effect of current pharmacological interventions for ADHD and provides neurobiological support for future clinical research focusing on reducing periphery-to-hub noise amplitude ratio and peripheral noise heteroscedasticity using targeted interventions including brain stimulation.

## Acknowledgments

This work was supported by the Ministry of Technology and Science, Taiwan (MOST103-2314-B-002-021-MY3), the National Health Research Institutes, Taiwan (NHRI-EX103-10008PI), and National Taiwan University Hospital (NTUH103-S2458, NTUH104-S2761). L.C. and J.A.R. are supported by the Australian National Health Medical Research Council (L.C., 1099082 and 1138711; J.A.R., 1145168 and 1144936).

## Supplementary Methods

### Participants and recruitment procedure

Recruitment occurred via advertisements at hospitals, colleges, and online. Potential adult participants were screened using the Taiwanese version of the Adult ADHD Self-Report Scale (ASRS) v1.1 [1]. Individuals deemed eligible to enter the study (i.e., they exhibited clinically relevant ADHD symptoms based on ASRS) were invited to the special clinic for adult ADHD at the Department of Psychiatry, National Taiwan University Hospital, Taipei, Taiwan for a clinical interview conducted by a board-certified child psychiatrist with extensive experience in ADHD diagnosis, intervention, and research across lifespan (author S.S.G.). Participants in the adult ADHD group were required to fulfil two criteria: i) they needed to currently demonstrate more than 6 items of ADHD symptoms as defined in the DSM-IV-TR in either inattentive, hyperactive-impulsive, or both domains; ii) ADHD symptoms must have occurred, or be noted before twelve years of age. This diagnosis of childhood-onset adult ADHD was based on both the clinical interview with the participants, and the Chinese version of the Kiddie-Schedule for Affective Disorders and Schizophrenia-Epidemiological version (K-SADS-E) interview with the parents. The ADHD diagnosis was further confirmed by the Conners’ Adult ADHD Diagnostic Interview (CAADI) [2] and the modified adult version of the ADHD supplement of the Chinese version of the K-SADS-E for childhood and current ADHD [3]. Besides ecologically-valid unstructured clinical interviews, DSM-IV psychiatric diagnoses were also confirmed by the semi-structured Chinese version of the Schedule of Affective Disorders and Schizophrenia-Lifetime (SADS-L) [3,4]. Matched healthy controls were recruited using the same procedure adopted for the ADHD group; Control participants received the same clinical evaluation and standard psychiatric interviews by S.S.G (i.e., the Chinese version of the K-SADS-E for childhood diagnoses, the CAADI, and the adult ADHD supplement plus SADS-L for ADHD and other psychiatric diagnoses at adulthood [5]). For both ADHD and controls, the following exclusion criteria were adopted: medical and mental illness other than ADHD, substance abuse, past or current use of psychotropic medication, and cognitive deficits (< 80 full-scale IQ measured by the Wechsler Adult Intellectual Scale-Third Edition [6]). Our sample selection strategy allowed the unequivocal assessment of brain mechanisms underpinning core symptoms of chronic ADHD [7]. However, the interpretation of current results must acknowledge that the specificity of our clinical group is biased and reduces generalizability (see main text).

### Measures for the validity of clinical diagnosis

The Chinese version of the K-SADS-E based on the DSM-IV-TR was developed by Gau et al. [5]. Rigorous methodological processes for the development of this instrument were implemented, including translation, back-translation, cultural adaptation, and assessment of psychometric properties.[5] To obtain information on ADHD symptoms and diagnoses in both childhood and adulthood, S. S. Gau further established a modified adult version of the ADHD supplement, which includes ADHD, oppositional defiant disorder, and conduct disorder derived from the Chinese K-SADS-E [8]. This adult version of the ADHD supplement and the Chinese SADSL have been widely used in our previous studies [3,4,9–13].

The study was approved by the Research Ethics Committee of the NTUH (201401024RINC) and registered as a clinical trial (NCT02642068). Written informed consent was obtained from all participants. Participants were recruited from March 2014 to December 2016.

### Measures of ADHD symptoms

The participants’ ADHD symptoms were dimensionally estimated by parent reports on the Chinese version of the Swanson, Nolan, and Pelham, version IV (SNAP-IV-C) scale [14] and self-reports on the Chinese version of the Adult ADHD Self-Report Scale (ASRS-C) [1]. The SNAP-IV-C, a 26-item scale, consists of Inattention (Item 1-9) and Hyperactivity/Impulsivity (Item 10-18), and Oppositionality (Item 19-26), corresponding to the core symptoms of ADHD and ODD on *DSM-IV-TR*, respectively. The 26 items of the SNAP-IV are rated on a 4-point Likert scale, with scores of 0-4 representing: “not at all,” “just a little,” “quite a bit,” and “very much.” The norms and psychometric properties of the Chinese version of the SNAP-IV (SNAP-IV-C) for parent reports have been established [14] and widely used in clinical and epidemiological studies in Taiwan. The raw scores of items 1-18 were used to measure the inattention and hyperactivity-impulsivity symptoms based on parents’ rating in the study.

The ASRS is a validated 18-question scale that was developed in conjunction with the revision of the World Health Organization Composite International Diagnostic Interview. The ASRS includes questions about the 9 inattention and 9 hyperactivity-impulsivity Criterion A symptoms of ADHD in the *DSM-IV-TR*, each question asking respondents how often a given symptom occurs over the past six months on a 0-4 scale (“never”, “rarely”, “sometimes”, “often”, “very often”). The psychometric properties of the Chinese version of the ASRS (ASRS-C) had been established in a sample of 4,329 Taiwanese young adults [1]. The ASRS-C has been widely used in adult ADHD studies in Taiwan.

### Imaging acquisition

The imaging protocol included: localizer, resting-state fMRI (7 min and 39 seconds), T1-weighted, and DSI. Functional images were acquired using a multi-echo EPI sequence: TR = 2.55 s; flip angle = 90°; matrix size = 64 × 64; in-plane resolution = 3.75 mm; FOV = 240 mm; 31 oblique slices, alternating slice acquisition slice thickness 3.75 mm with 10% gap; iPAT factor = 3; band-width = 1698 Hz/pixel; echo time, TE = 12, 28, 44 and 60 msec). T1 images were acquired using an MPRAGE sequence with a TR = 2 s; TE = 2.98 msec; flip angle = 9°; matrix size = 256 × 256; inversion time = 900 msec; voxel size = 1 mm^3^. DSI acquisitions used a pulsed-gradient spin-echo diffusion EPI sequence with a twice-refocused balanced echo repetition time/echo time = 9600/130 msec, slice thickness = 2.5 mm, acquisition matrix = 80 × 80, field of view = 200 × 200 mm, in-plane spatial resolution = 2.5 mm × 2.5 mm, 101 diffusion-encoding directions covering a half q-space 3D grid with radial grid size of 3, b_max_= 4000 s/mm^2^ [15].

### MRI preprocessing

In short, the resting-state fMRI preprocessing pipeline included: quality control, comprehensive data denoising using multi-echo independent components analysis (ME-ICA v3.0)[34], coregistration to individual anatomical images, non-linear normalization to MNI space, and filtering (0.01∼0.1 Hz). The full preprocessing pipeline is reported elsewhere [9].

The DSI data underwent an initial quality assurance procedure: Individual DSI images [54 slices × (101 directions DW images + 1 null image) = 5,508 images] were scrutinized by calculating signals in the central square (20 × 20 pixels) of each image. Signal loss was defined as the average signal intensity of an image lower than two standard deviations from the mean of all images (after correcting for its b value) [16]. As jerky head motion induces a signal loss in DSI images, these signal dropout counts were considered a proxy estimate for overall levels of in-scanner head motions. Individuals’ DSI data with more than 90 images of signal loss, at either baseline or follow-up, were excluded from further analyses [16].

DSI data were reconstructed using the *q*-space diffeomorphic reconstruction (QSDR) approach implemented in DSI Studio (www.dsi-studio.labsolver.org) [17]. QSDR first computed the quantitative anisotropy in each voxel in native space. Then the reconstructed images were warped to a template in Montreal Neurological Institute (MNI) space using constrained diffeomorphic mapping. In MNI space, a diffusion sampling length ratio of 1.25 mm with five fiber orientations per voxel and 8-fold orientation distribution function tessellation (642 sampling directions) was used to obtain the spin distribution function, and the output resolution was 2 mm. A deterministic fiber tracking algorithm [18] was performed with extreme turning angle threshold of 60°, step size of 1.0□mm, minimum and maximum lengths of 10LJ and 400 mm, respectively. 10,000,000 streamlines were seeded throughout the whole brain and terminated when the local quantitative anisotropy fell below values estimated using Otsu’s threshold [18], which gives the optimal separation between background and foreground. Other tracking parameters as specified in DSI Studio were: smoothing: 0; seed orientation: all; seed position: subvoxel; randomize seeding: off; direction interpolation: trilinear.

### Head motion

Micro-head movements (mean framewise displacement, FD) [19] for rs-fMRI and signal dropout counts [16] for DSI, were not significantly different between ADHD and controls (p = 0.35 and 0.54, respectively). Individual differences in subject head motion during structural and functional data acquisition did not correlate with ADHD symptoms (first principal component, **Supplementary Table 3**); DSI motion-related signal loss: r = −0.02, p = 0.84 in ADHD, r = −0.16, p = 0.16 in controls; resting state FD: r = 0.14, p = 0.13 in ADHD, r = 0.08, p = 0.50 in controls (see **Figure S1**).

**Figure S1.**
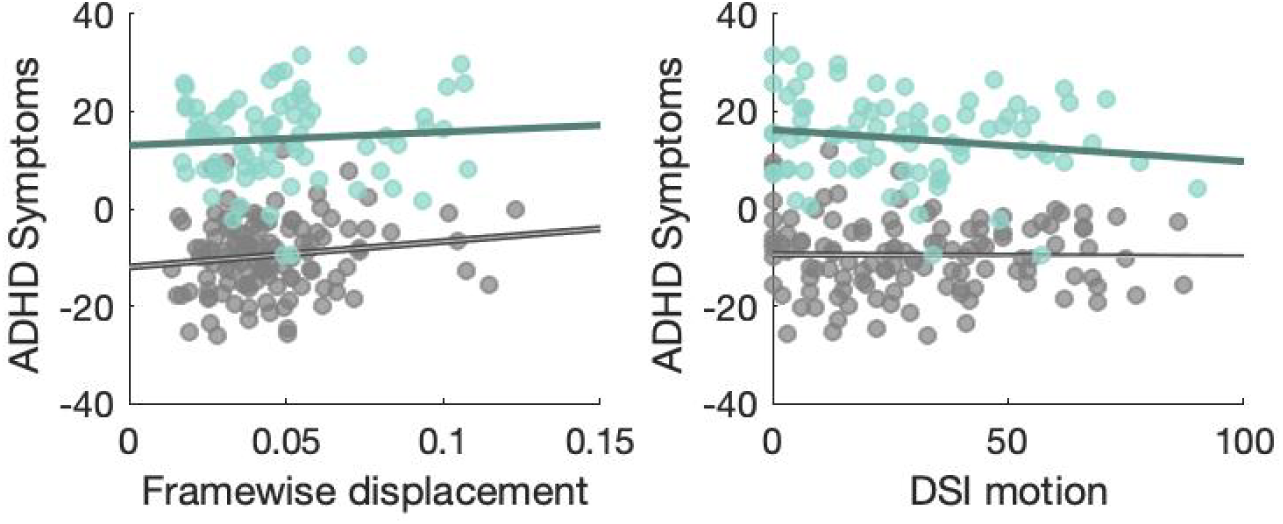
Correlation between functional (framewise displacement, left) and structural (DSI motion, right) head motion and ADHD symptoms. Neither ADHD (teal colors) or control (grey colors) demonstrated significant associations between head motion and symptoms.

Varying definitions of brain network hubs exist [20]. Here, we identified hub-regions according to aggregate ranking across multiple metrics [21,22]. First, for each participant, each node’s “hubness” was calculated from its composite average ranking across degree, strength, betweenness and subgraph centrality scores using the brain connectivity toolbox [23]. The top 15% composite scores (N = 32, **Figure S2, Supplementary Table 1&2**) were used to identify hub-regions within each participant; all other nodes were assigned as *periphery* nodes.

**Figure S2.**
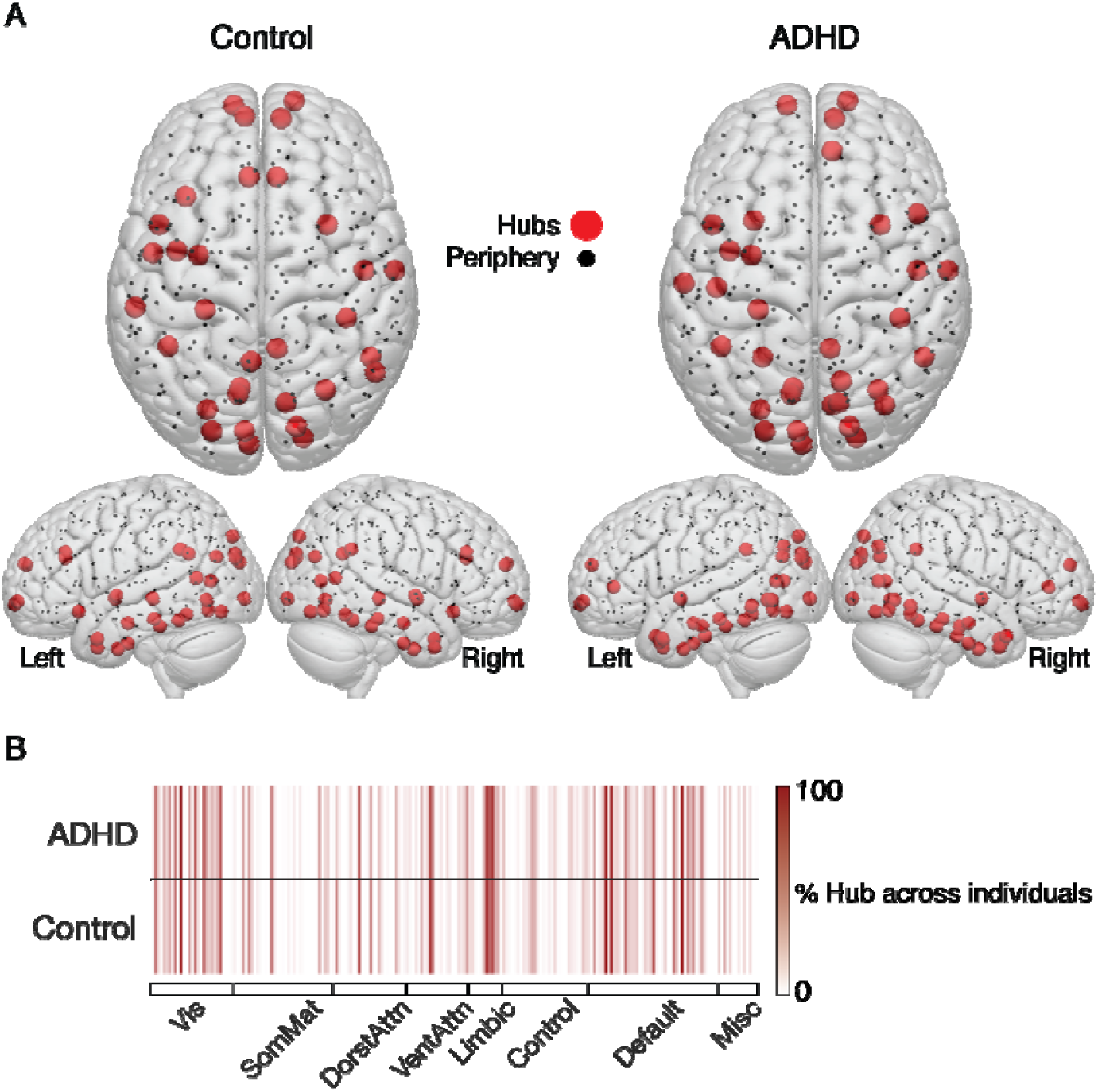
Structural hub topology in adult ADHD and healthy matched controls. **A.** Brain rendering of group average hub (red) and periphery (black) nodes. **B.** Individual-level representation of hub regions in canonical resting-state brain networks. Darker lines indicate more consistency within each group (i.e., dark red represents every individual had a hub node within the ADHD or Control group). Overall, this plot highlights the consistency in detecting hub regions at the level of single subjects between groups.

### Statistical comparisons between groups

Across all statistical tests within-group data distributions were not normal (Kolmogorov-Smirnov test, p < 0.05), thus non parametric statistics were used (the Mann–Whitney U test). A significant Mann-Whitney U test can be interpreted as either a difference in distribution, or a difference in the medians between two groups. Thus for significant findings we evaluated whether the variances between groups were similar. Levene’s test (p > 0.05) confirmed that the variance between the groups was similar, which suggests a difference in medians across groups.

### Empirical results control analyses

A number of tests were conducted to establish the reliability of our empirical findings. To ensure that our chosen brain parcellation had little bearing on the results [24], we repeated the analyses in two other, independent brain parcellations: *Shen-213* [25] and *Brainnetome-244* [26]. The reported effects were all successfully replicated (**Supplementary Table 2**). Using these alternative brain parcellations, we also found that adults with ADHD exhibited weaker structure-function coupling in hub connections. However, the effect size of these between-group differences was consistently smaller than the effect in feeder connections.

### Computational modeling

The dynamics of each region in the connectome are modeled as a multivariate Ornstein-Uhlenbeck process with independent white Gaussian noise drive, and obey the following stochastic differential equation:

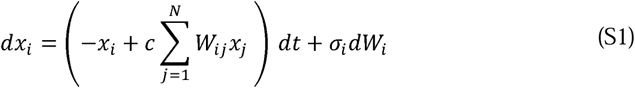

where *x*_*i*_ is the activity of the *i*-th region; *c* is the global coupling strength which rescales the strength of structural connections of the system; *W*_*ij*_ is the connectivity weight to region *I* from region *j* (as specified by the empirical SC matrix); *σ*_*i*_ is the intrinsic noise amplitude/level of the *i*-th region, and defines the size of zero-mean Gaussian random increments increments/steps in the dynamics of the region, and *N* is the total number of regions in the connectome.

In this work, we hypothesized that each region has a different value of. **Figure S3** shows schematics of how the heterogeneous intrinsic noise levels, in combination with the interactions defined by the SC matrix, affect the functional timeseries of a subject. **Figure S3A** illustrates probability density functions (pdfs) of the random increments in Gaussian noise () scaled by in Eq. (S1), for three different regions colored in red, blue and yellow. The mean of each pdf is assumed to be zero, while the standard deviation of each pdf is, i.e., the intrinsic noise amplitude of the region. **Figure S3B** shows the expected functional timeseries of the same regions illustrated in **Figure S3A**. The second-order fluctuations (i.e., variance) of these timeseries are related to the intrinsic noise amplitude (). While estimating empirically is possible [27], longer recording sessions would be needed to derive these values accurately for each individual [28]. Thus, we tested a relatively broad range and distributions for that preserved the validity of the model. It is also important to note that the current model produces a 1/f^α^ noise spectrum as detected in emprirical EEG data (**Figure S4**).

**Figure S3.**
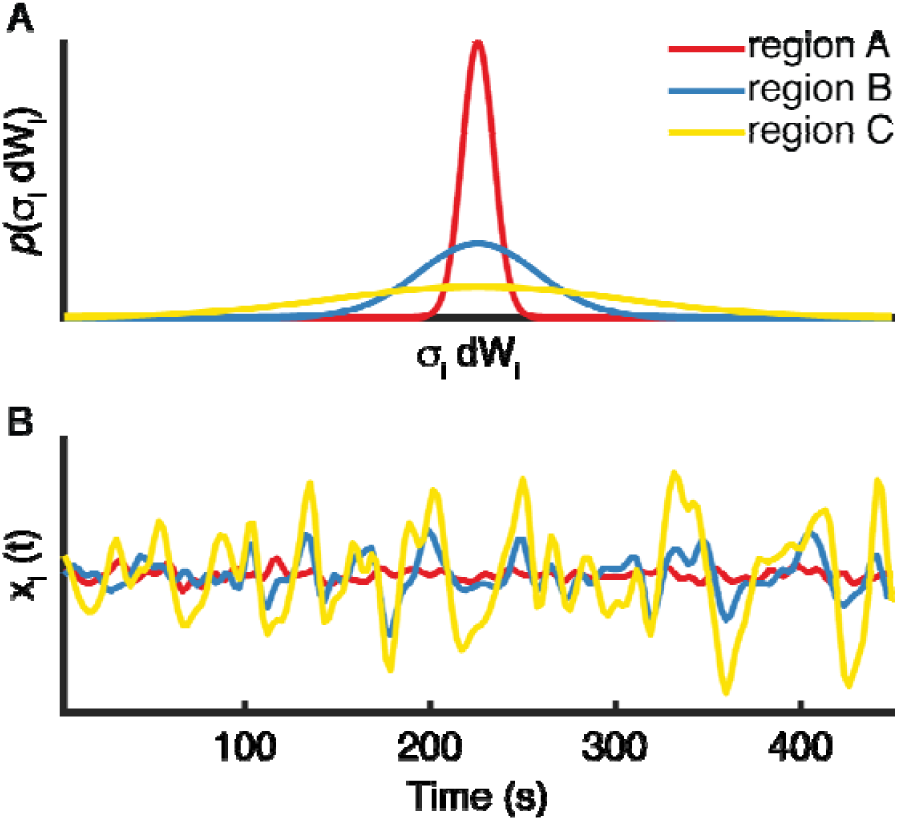
Schematic of functional time series with different intrinsic noise amplitudes (σ_i_). **A.** The probability density distribution *p*(*σ*_*i*_*dW*_*i*_) of the random increments in Gaussian noise *σ*_*i*_*dW*_*i*_ for three regions A (red), B (blue) and C (yellow). The values of *σ*_*i*_ increase in the following order *σ*_*A*_ < *σ*_*B*_ < *σ*_*C*_. That is, region A has the smallest noise amplitude whereas region A has the largest. **B.** Exemplar modeled functional timeseries for regions A (red), B (blue) and C (yellow), showing the effect of increasing *σ*_*i*_.

**Figure S4.**
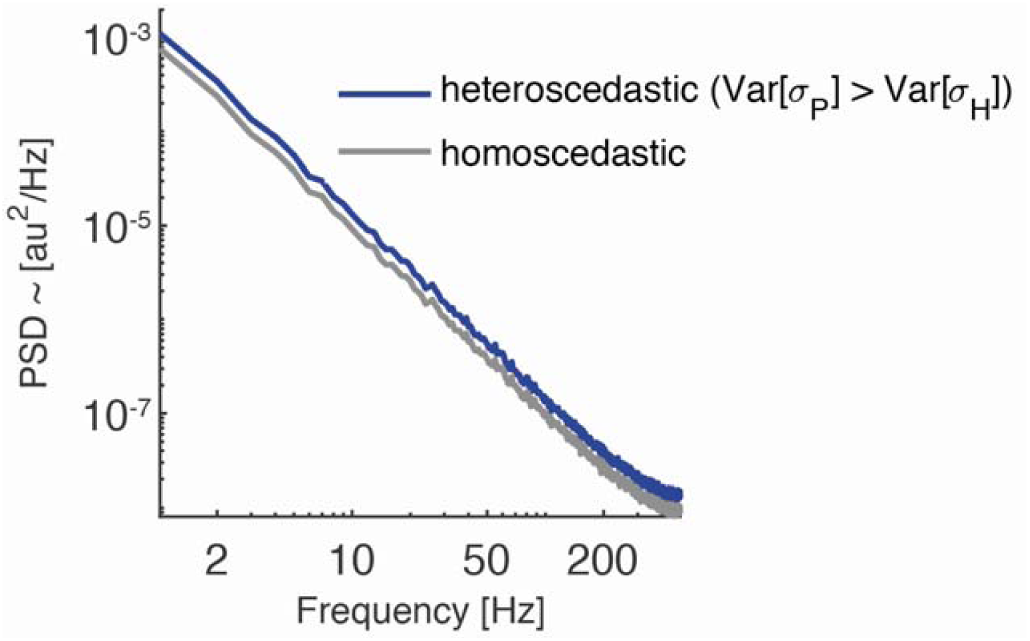
Power spectra of neural noise produced by our model. The spectra follow the characteristic 1/f^α^ law as detected in empirical EEG data. The power spectral density (PSD) is expressed in arbitrary units (a.u ^2^/Hz). Gray line: Homoscedastic case E[σ_P_]=E[σ_H_]=1, and Var[σ_P_]=Var[σ_H_]=0. Green line: Heteroscedastic case with E[σ_P_]>E[σ_H_] (1.5 and 1.2, respectively), and Var[σ_P_]>Var[σ_H_] (0.2 and 0.02, respectively). Compared to the homoscedastic case, the heteroscedastic case is characterized by increases in neural noise. This is reflected by the curve shifting upwards toward higher power.

### Effects of heteroscedasticity on the structure-function coupling

We hypothesize that increased *heteroscedasticity*, that is, increased heterogeneity in the intrinsic noise levels across brain regions is a driving factor for the observed breakdown between structure and function in ADHD. Thus, to systematically study the effects of *heteroscedasticity* we use the model described in the previous subsection. To select the values of *σ*_*i*_, we first consider the fact that there are two main groups of regions: hub regions (*H*) and peripheral regions (*P*). The individual intrinsic noise amplitudes (*σ*_*i*_) are assumed to be samples of either *σ*_*H*_ or *σ*_*P*_, which are random variables with distinct expected (mean) values (E[·]) and variances (Var[·]): (E[*σ*_*H*_], Var[*σ*_*H*_]) and (E[*σ*_*P*_], Var [*σ*_*P*_]).

The first scenario analyzes the effect of distinct expected values in hubs and periphery regions (E[*σ*_*H*_] *vs.* E[*σ*_*P*_]), while assuming Var[*σ*_*H*_] = Var[*σ*_*P*_] = 0. This study produced the 2D maps presented in **Figure 3A-C** in the main text. These maps were calculated as the average of 118 individual maps using the SC matrices of control subjects. On the other hand, the 2D maps of the second study presented in **Figure 3D-F** (main text), calculated as the average of 1888 individual maps, one per SC matrix of the control group (118 subjects); and, 16 different seeds of the pseudorandom number generator used to draw the values of individual σ*i* for each combination of (Var[*σ*_*H*_], Var[*σ*_*P*_]).

The second scenario analyzes the effects of varying the noise levels between the groups and also from region to region (E[*σ*_*H*_] ≠ E[*σ*_*P*_] and Var[*σ*_*H*_]≠ Var[*σ*_*P*_])). Specifically, we calculated 2D maps of SC-FC coupling for hub, feeder, and local connections for varying degrees of intra-hub and intra-periphery heteroscedasticity, for different values of asymmetry between E[*σ*_*H*_] and E[*σ*_*P*_]: 0%, 10%, 20% and 50%, always for *σ*_*H*_ < *σ*_*P*_. The 2D maps of SC-FC coupling for feeder connections are shown in **Figure S5 A-D**. The bottom panels (**E-G**) are the ‘percentage difference maps’ between the 2D map in **Figure S5 A** with E[*σ*_*H*_]= E[*σ*_*H*_], and the maps with an asymmetry between mean noise levels in hub and periphery regions (**Figure S5 B-D**). Decreases in SC-FC coupling with respect to the map shown in **Figure S5 A** are shown in blue, while increases are shown in red. These results show that: (i) the presence of an asymmetry in the noise level between groups is the primary driving factor of SC-FC decoupling, illustrated by the shrinking area of high SC-FC coupling in **Figure S5 A-D**; and, (ii) increased heteroscedasticity within the peripheral regions is the second factor that further breaks down the SC-FC coupling, illustrated by the blue areas along the direction of increasing Var[*σ*_*P*_].

**Figure S5.**
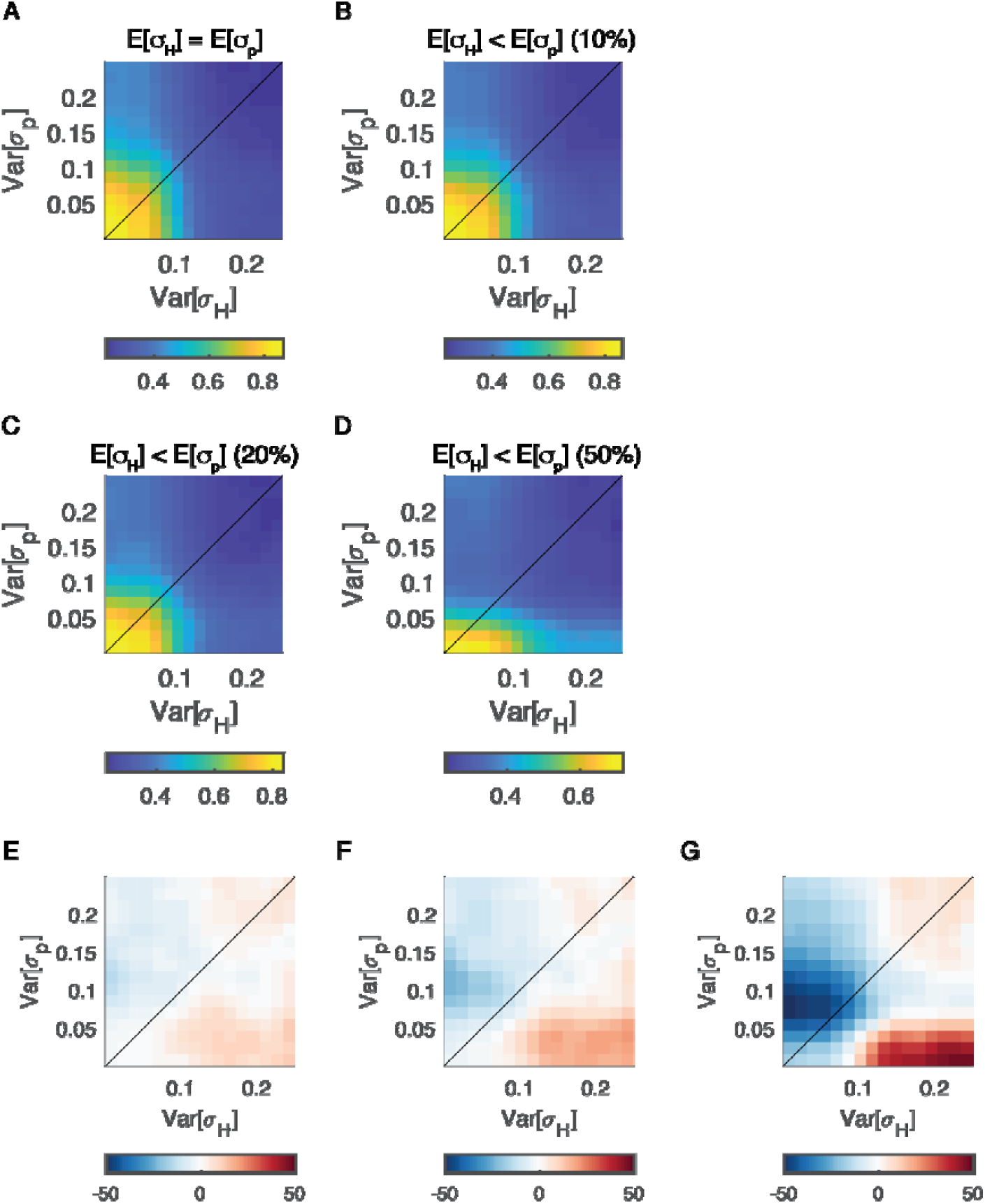
Maps of structural-functional connectivity coupling using different levels of within- and between-group noise heteroscedasticity (top row). The bottom row shows maps of the percentage difference between the corresponding panels above and the null between-group heteroscedasticity in panel A. **A.** Between-group noise heteroscedasticity is 0 (=). **B.**is 10% larger than. **C.**is 20% larger than. **D.**is 50% larger than. **E.** Percentage difference map between map in panel A and B. **F.** Percentage difference map between map in panel A and C. **G.** Percentage difference map between map in panel A and D. The colors represent absolute SC-FC coupling (top row), and positive (red) and negative (blue) percentage difference with respect to the SC-FC couplings in panel A (bottom row).

## Supplementary Tables

**Supplementary Table 1.**
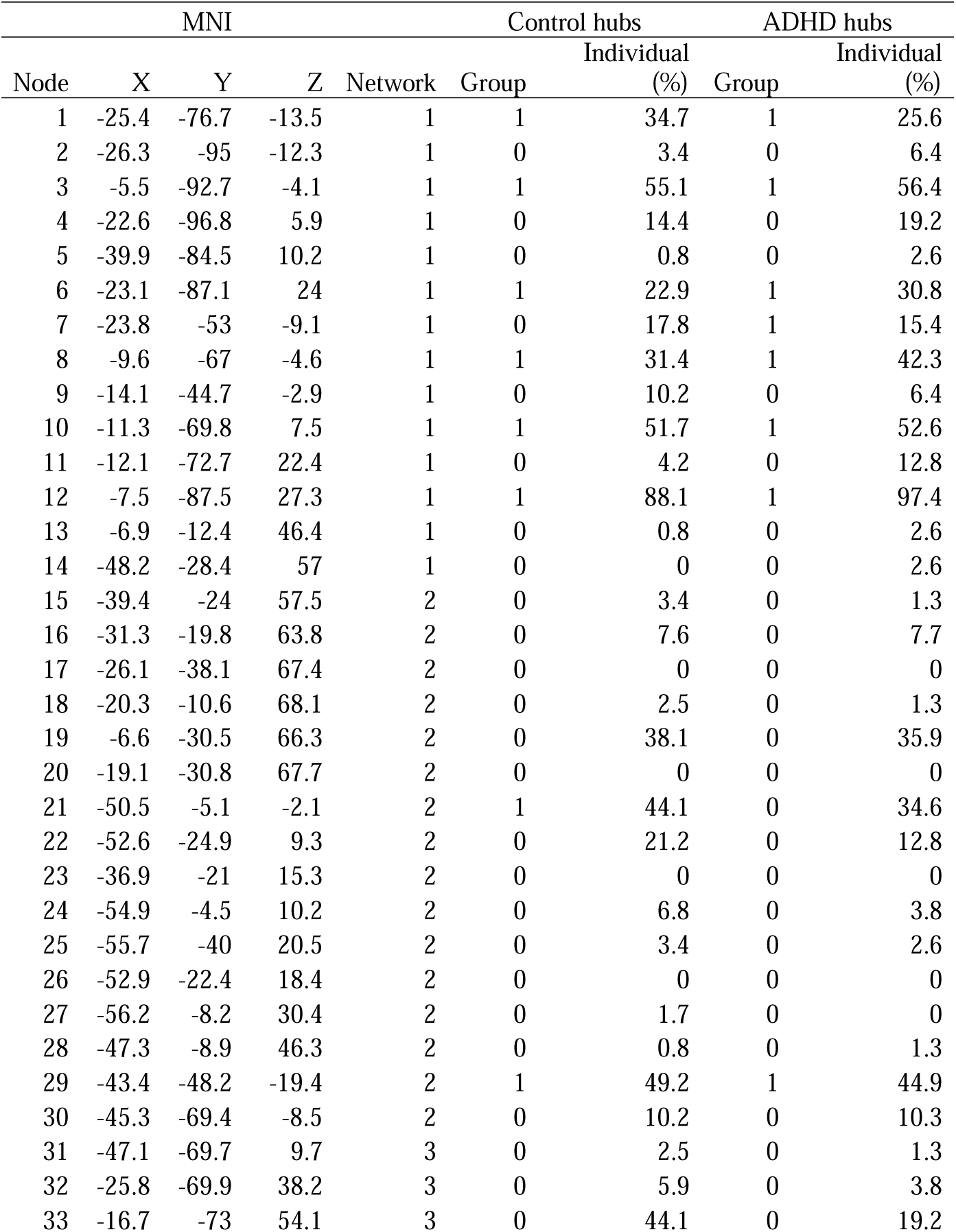
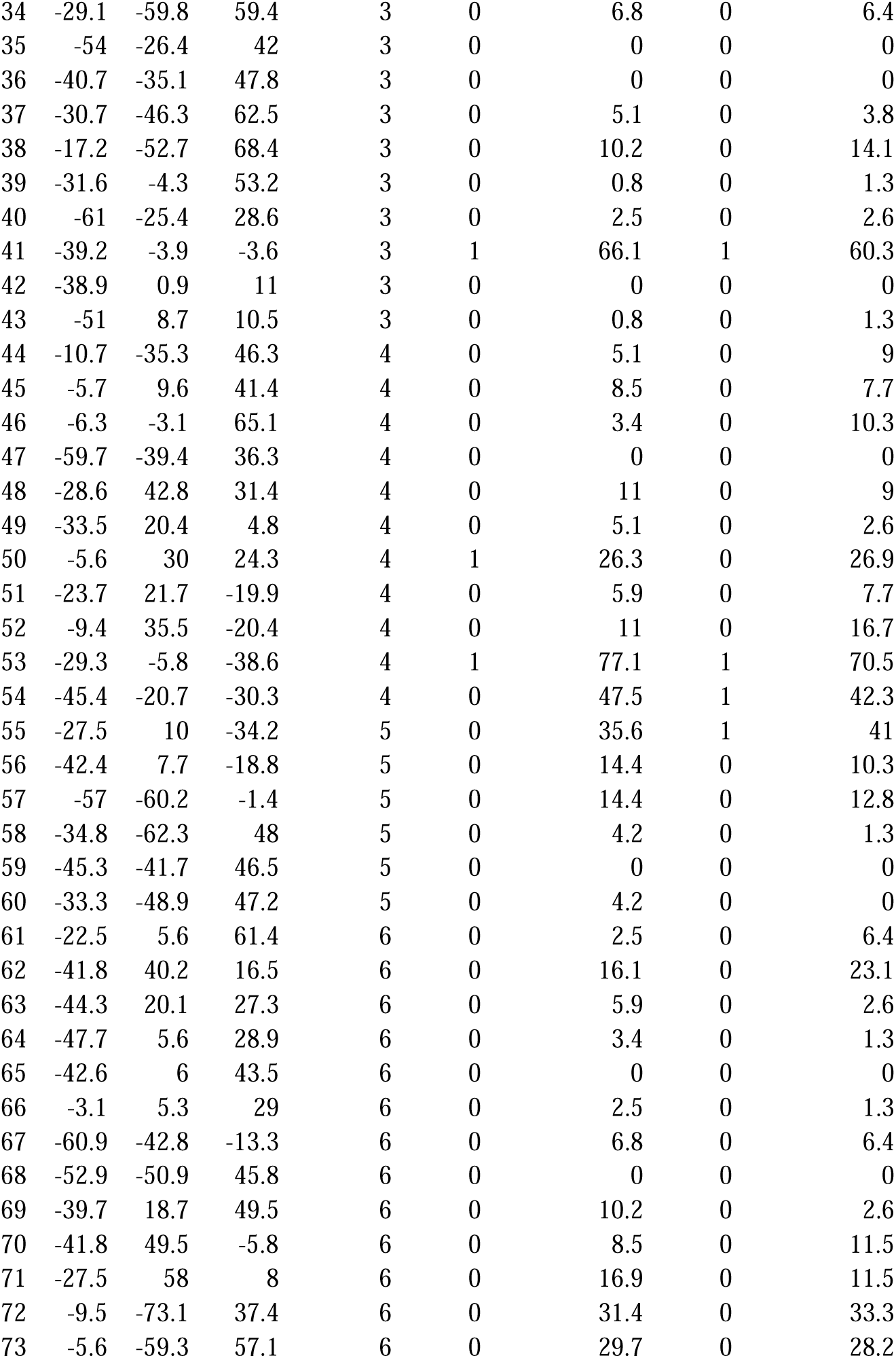
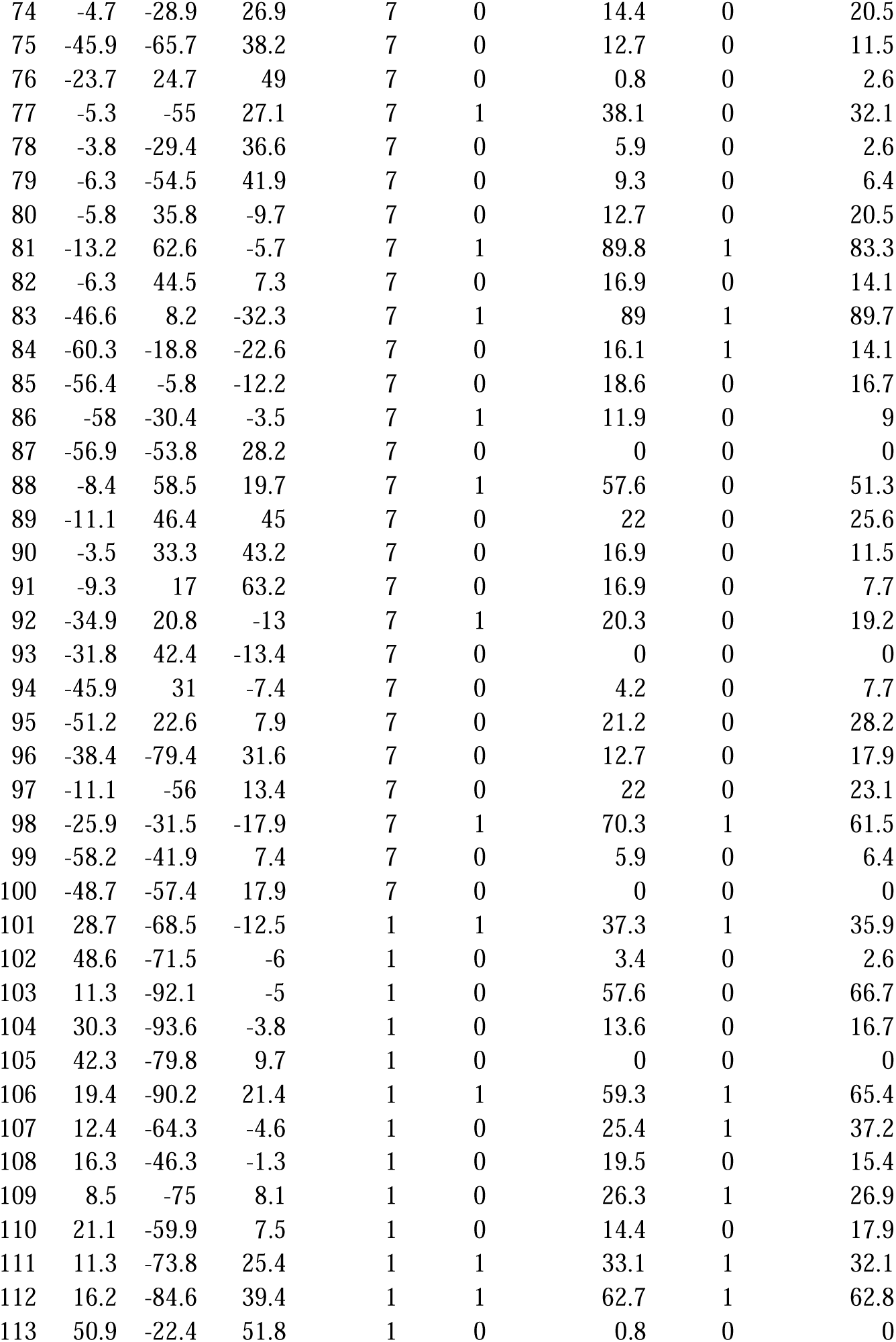
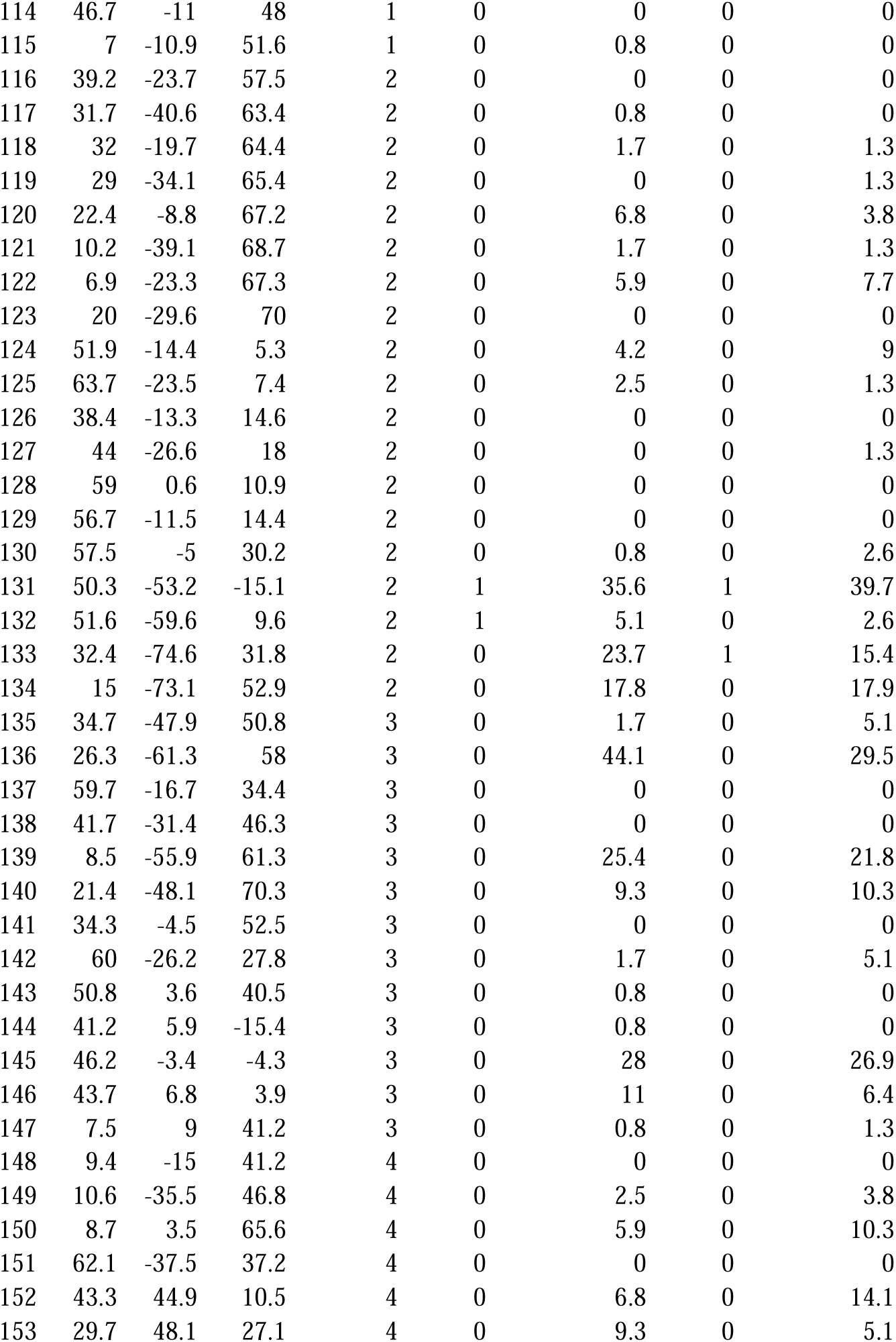
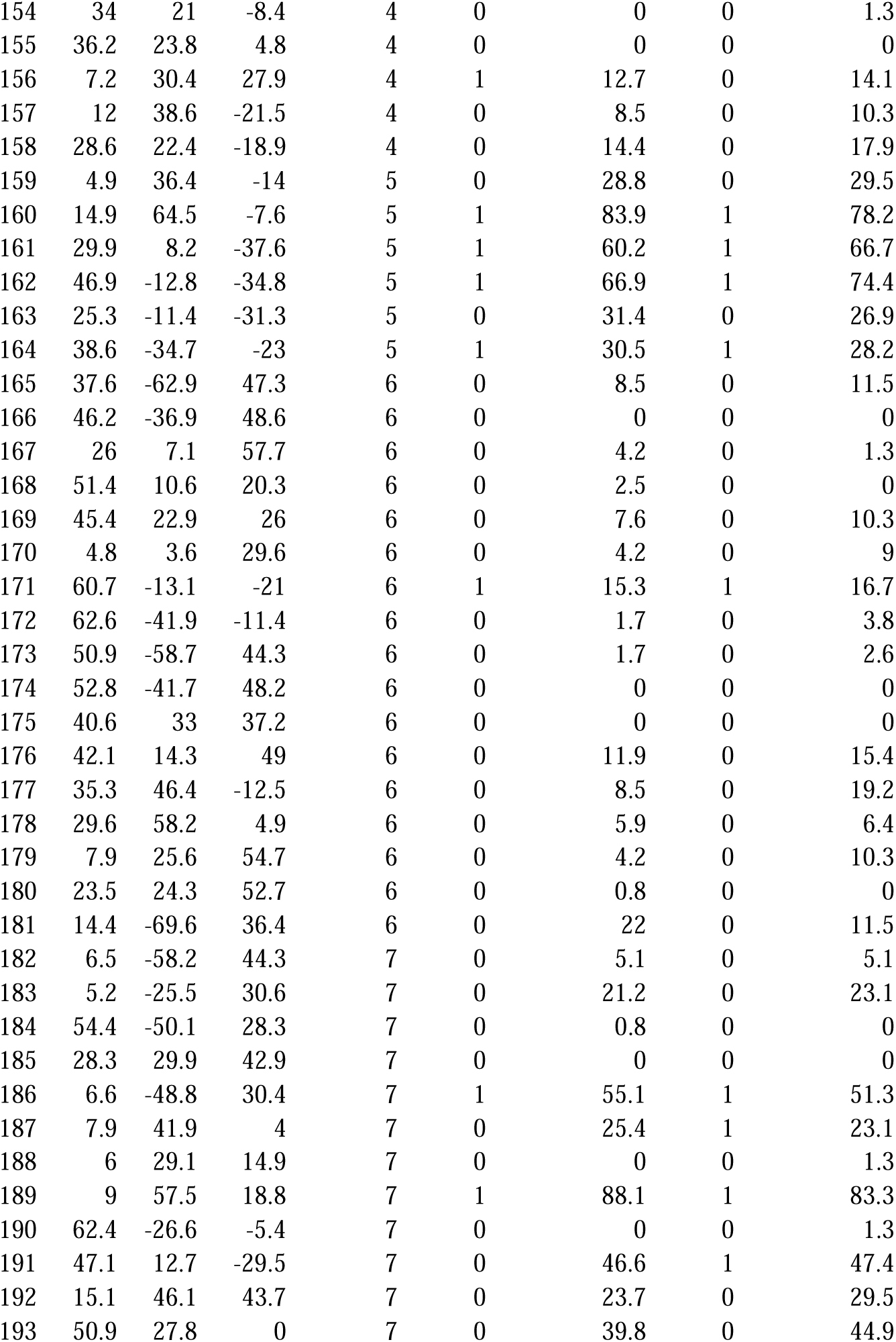
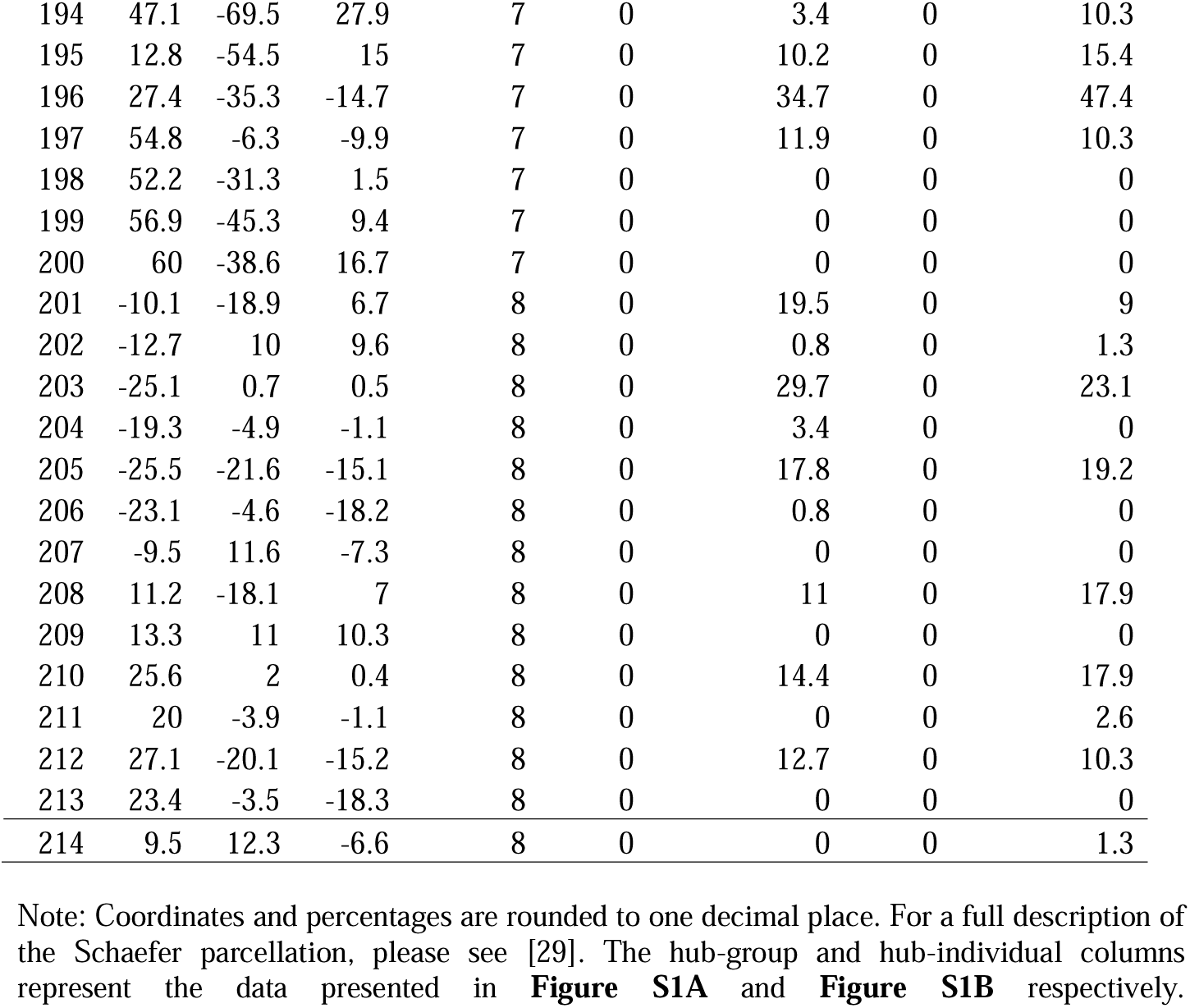
Table of Schaefer 214 coordinates, network assignments, hub status for CTRL and ADHD

**Supplementary Table 2.**
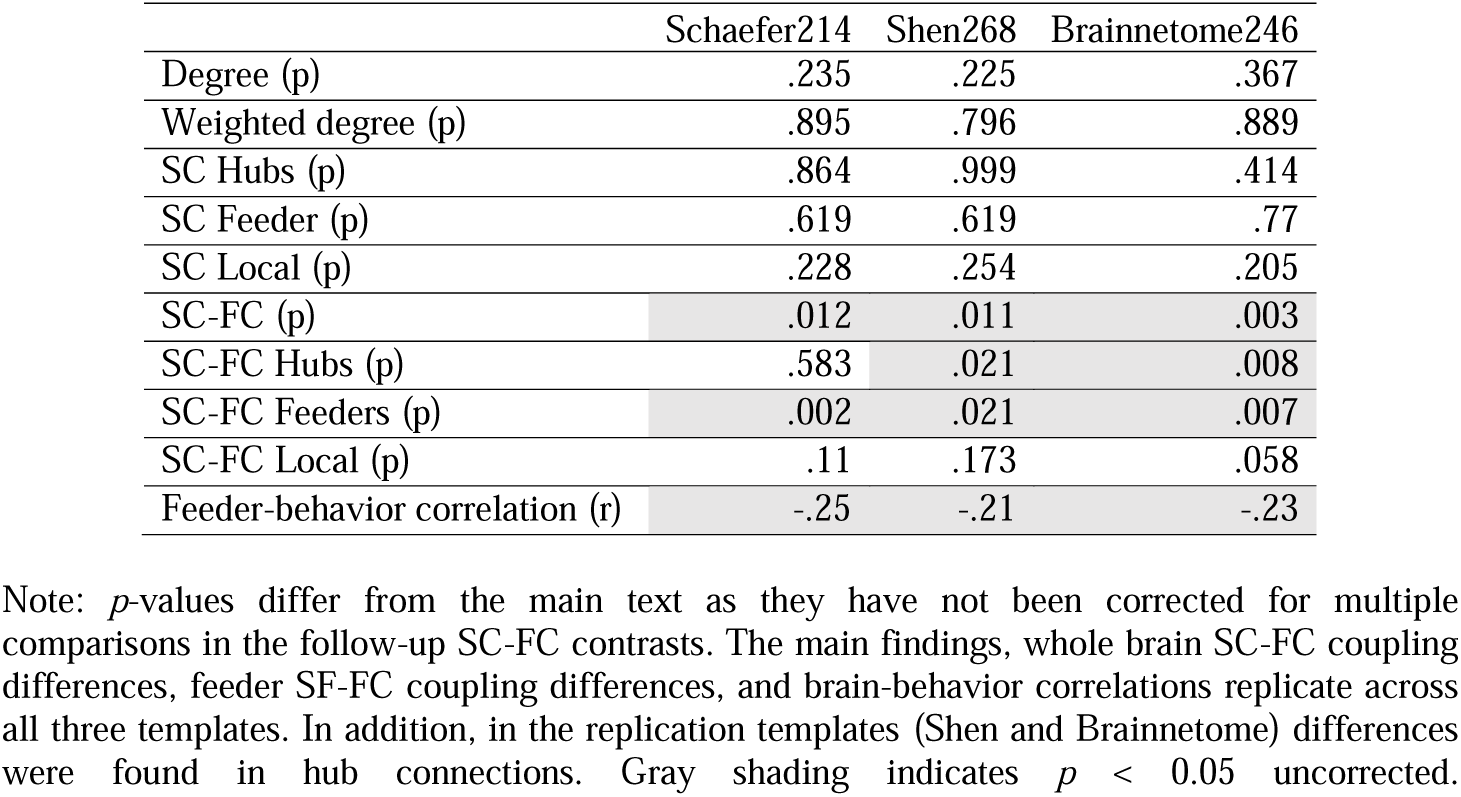
Empirical comparison across brain parcellations

**Supplementary Table 3.**
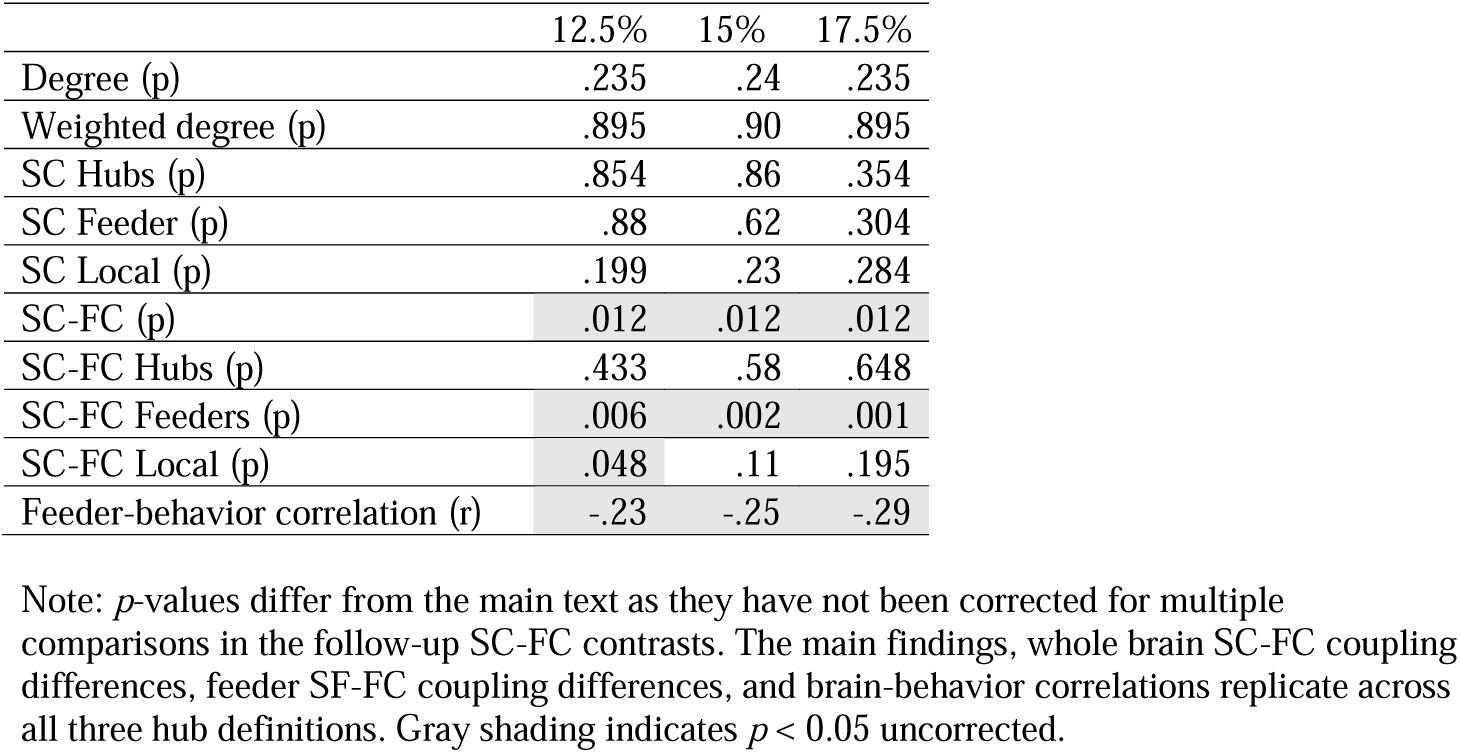
Effect of hub definition on results

**Supplementary Table 4.** Principal component analysis loadings

|  | Component1 | Component2 | Component3 | Component4 |
| --- | --- | --- | --- | --- |
| Inattention SNAP-IV (parent-rated) | .50 | -.5 | -.47 | -.53 |
| Hyperactivity/Impulsivity SNAP-IV (parent-rated) | .48 | -.52 | .56 | .44 |
| Inattention ASRS (self-rated) | .52 | .41 | -.5 | .56 |
| Hyperactivity/Impulsivity ASRS (self-rated) | .50 | .56 | .46 | -.46 |
| **Variance explained** | **80.72** | **9.49** | **6.68** | **3.11** |

